# Use of whole-genome sequencing in the epidemiology of *Campylobacter jejuni* infections: state-of-knowledge

**DOI:** 10.1101/078550

**Authors:** Ann-Katrin Llarena, Mirko Rossi

## Abstract

High-throughput whole-genome sequencing (WGS) is a revolutionary tool in public health microbiology and is gradually substituting classical typing methods in surveillance of infectious diseases. In combination with epidemiological methods, WGS is able to identify both sources and transmission-pathways during disease outbreak investigations. This review provides the current state of knowledge on the application of WGS in the epidemiology of *Campylobacter jejuni*, the leading cause of bacterial gastroenteritis in the European Union. We describe how WGS has improved surveillance and outbreak detection of *C. jejuni* infections and how WGS has increased our understanding of the evolutionary and epidemiological dynamics of this pathogen. However, the full implementation of this methodology in real-time is still hampered by a few hurdles. The limited insight into the genetic diversity of different lineages of *C. jejuni* impedes the validity of assumed genetic relationships. Furthermore, efforts are needed to reach a consensus on which analytic pipeline to use and how to define the strains cut-off value for epidemiological association while taking the needs and realities of public health microbiology in consideration. Even so, we claim that ample evidence is available to support the benefit of integrating WGS in the monitoring of *C. jejuni* infections and outbreak investigations.

**Abbreviations**
CC: clonal complex
cgMLST: core-genome MLST
DALY: Disability-adjusted life year
ECDC: European Centre for Disease Prevention and Control
EFSA: European Food Safety Authority
Epi-linked: epidemiologically linked
GWAS: genome-wide association study
MALDI-TOF: Matrix Associated Laser Desorption Ionization–Time of Flight
MLST: multilocus sequence typing
NGS: Next generation sequencing
SNPs: single nucleotide polymorphisms
SNV: single nucleotide variant
ST: Sequence type
PFGE: Pulsed-field gel electrophoresis
wgMLST: whole-genome MLST
WGS: Whole-genome sequencing

## INTRODUCTION

The ability to conduct epidemiological investigations and intervene to control and prevent food- and environmentally transmitted diseases is a major task of public health authorities. In this regard, identifying epidemiologically (epi)-linked cases and differentiating them from concurrent, sporadic incidences are essential in risk assessment, outbreak investigations, and source attribution. To do so, traditional epidemiological investigations is concurrently used with molecular subtyping of the etiological agent, and no other method offers a higher degree of resolution than whole-genome sequencing (WGS). The recent development of high throughput sequencing technologies for WGS (next-generation sequencing, NGS) has prompted large-scale detailed studies of entire pathogen genomes, replacing the need for traditional typing methods [e.g. pulsed-field gel electrophoresis (PFGE), serotyping, biotyping] and sequence-based investigations (e.g. resistance or virulence gene detection). Moreover, the declining costs of NGS and availability of bench-top analyzers facilitate the application of WGS in routine surveillance and outbreak investigations of bacterial and viral infectious disease by public health authorities (1). As a result, WGS analysis is currently used in real-time surveillance of *Listeria monocytogenes* and *Salmonella* Enteritidis by the American Centers for Disease Control and Prevention and the U.S Food and Drug Administration (http://www.fda.gov/Food) and such approaches for other foodborne pathogens are expected to come in use shortly. Furthermore, both the European Food Safety Authority (EFSA) and the European Centre for Disease Prevention and Control (ECDC) have stated that WGS should be the analysis method of choice in public health activities (2,3).

*Campylobacter jejuni* is the one of the most frequent causes of bacterial gastroenteritis globally, with an estimated burden of illness of 7.5 Disability-adjusted life year (DALY) (4,5). The epidemiology of *C. jejuni* is only partially understood and shedding new light on this area is difficult as most cases appear to be sporadic and remain unreported (6). Additionally, the occurrence of immune individuals in the population coupled with the many possible reservoirs, transmission pathways, and regional epidemiological differences makes source tracing and risk assessment difficult (7). Therefore, more knowledge on these matters is needed to curb the current campylobacteriosis epidemic. Until now, the two most common molecular typing methods [PFGE and the seven-loci multilocus sequence typing (MLST) scheme], have been irreplaceable working tools in studying the epidemiology of *C. jejuni* infections, and have greatly contributed to the current knowledge on the nature of *C. jejuni* infections in human patients and their potential reservoirs (8–19). However, limitations of these methods has been uncovered which deem PFGE and MLST unsuitable as sole subtyping methods for assessing epidemiological links between *C. jejuni* isolates (20–22). Therefore, developing an efficient “second generation” molecular-guided epidemiological surveillance is clearly necessary for *C. jejuni* and of high priority to overcome the limitations of available typing methods (23). As international standardization is needed due to the global and regional nature of campylobacteriosis, full comparability of molecular data of strains collected from different areas must be possible. Great hopes have been placed on WGS approaches in these regards, as WGS offers a high discriminatory power that allows detection of all possible epidemiologically significant variation between strains (23–25) and the global and regional standardization and comparison achievable through WGS, which today is facilitated by public databases (26,27).

In this review we show how WGS has improved, or has the capacity to improve, surveillance and outbreak detection of *C. jejuni* infections and discuss how WGS has increased our understanding of both the evolutionary and epidemiological dynamics of campylobacteriosis caused by *C. jejuni*.

## METHODS

A scientific literature review was carried out by searching the Google Scholar database (scholar.google.com), using the qualifiers ‘Whole-genome sequencing AND *Campylobacter jejuni*’, and ‘epidemiology OR ecology’. Search was done in August-September 2016 resulting in a list of 2,640 publications. Inclusion criteria were based on title and abstract, leading, if related to the use of WGS in epidemiological investigation of campylobacteriosis, to retrieval and analysis of the full paper. Non-peer-reviewed studies and studies focused solely on functional genomics were not included. In addition, the authors integrated their own knowledge and opinions with the scientific literature review.

### SUBTYPING OF *C. JEJUNI* IN OUTBREAKS AND SURVEILLANCE; ADAPTING WGS TO THE MOLECULAR EPIDEMIOLOGY OF CAMPYLOBACTERIOSIS

An outbreak is the sudden increase in the number of expected cases of a disease in a population in a limited geographical area over a short period of time. Outbreak detection depends on knowledge of the baseline level of that disease, which is usually acquired through surveillance (28). Additionally, through data analysis and isolate collection from monitoring programs it is possible to detect so-called diffuse outbreaks, i.e. clusters of cases with a presumptive common source not necessarily clustered geographically and/or represented by an increase in cases above the expected baseline level (29). To be effective, surveillance depends on correct reporting of communicable diseases, and the EU has therefore provided a case definition and clinical and laboratory criteria for diagnosis of *C. jejuni* infections to secure and unify reporting of campylobacteriosis (30).

Upon outbreak detection, epidemiologists and microbiologists work together to trace the source and include or exclude patients as part of a given outbreak by combining traditional epidemiological tools and the analysis of samples collected from sources and patients. From a microbiologists’ point of view, isolation of the causative pathogen with subsequent subtyping of that agent is critical. As mentioned above, one of the subtyping method most commonly used in campylobacteriosis outbreaks investigations has been PFGE (16,17,19,31). PFGE is highly discriminatory when used with certain restriction enzymes (31), and the method of Tenover *et al.* (1995) is commonly applied to decide whether two isolates are related or not (32). However, PFGE can both over- and underestimate the clonal relationship between *C. jejuni* strains (20).

Additionally, MLST has been the essential subtyping method for longitudinal analysis of *C. jejuni* isolates acquired through monitoring programs (8,9,(11–15), but lacks the necessary resolution for cluster analysis as extensive genomic mosaicism between highly related strains has been demonstrated (33).

WGS is currently starting to take over as the typing method of choice for *C. jejuni* outbreaks investigation, and has the potential to substitute MLST in routine surveillance of campylobacteriosis. However, several issues regarding the epidemiology and genomic diversity need to be addressed before WGS can be a useful and reliable working tool in the public health sector. In particular, understanding baseline ecological diversity and population structure of *C. jejuni* and the genomic diversity within a single human infection is a necessary prerequisite for determining if isolates are clonal, as would be expected in a cluster of cases or outbreak with a common source. Moreover, the resolution depth of WGS should be adjusted to balance the power to resolve isolates from the main *C. jejuni* lineages while maintaining clonal stability between epi-linked strains from outbreaks and source and patient. The latter might include creating and agreeing on a suitable nomenclature system for *C. jejuni.* Here we review how laboratory diagnosis and surveillance of *C. jejuni* infections (species determination, subtyping and the definition of clonal isolates) are currently performed and how the implementation of WGS could change, or have already changed, this field of public health.

### Species determination

Determining the causative agent present in a sample of stool or food is crucial in making effective clinical decisions and public health interventions. Today, species determination is accomplished by culturing the pathogen on selective or non-selective media, identifying suspicious colonies, and producing a mass of pure organism for further use. For *C. jejuni*, growth on selective media containing antibiotics and blood or charcoal antioxidants in a microaerophilic atmosphere is the most common choice, and was instrumental in establishing *C. jejuni* as a major cause of human gastroenteritis in the late 1970s and early 1980s (34,35). Thereafter, colony morphology, Gram stain, motility-, oxidase- and hippurate hydrolysis tests aid to identify *C. jejuni* (36,37). Molecular methods, such as genus-specific PCRs detecting the *hipO-*gene (38) or sequencing the 16sRNA, *groEL* (39) or the *rpoB* (40) genes have been used to differentiate between *Campylobacter* species. The entire process can take approximately one to two weeks, which is a considerable amount of time when immediate intervention or treatment is needed.

The recent introduction of Matrix Associated Laser Desorption Ionization–Time of Flight (MALDI-TOF) mass spectrometry for rapid and cost effective species determination of bacterial pathogens (41,42) has considerably shortened the time needed for identification of *Campylobacter* species, especially in clinical settings. The method allows precise identification [99.4% accuracy for *Campylobacter jejuni* (42)] of bacterial colonies collected directly from the primary isolation plate. As MALDI-TOF mass spectrometry is faster and more accurate than standard methods, it will probably remain the reference methodology in the future, regardless of the WGS revolution.

Although the use of WGS would increase the confidence of species delineation in most cases, current detection methodologies for *Campylobacter*, coupled with MALDI-TOF, are generally cheap and fast enough to satisfy the need for both clinical diagnostic and surveillance. In this scenario, implementation of WGS for species determination would advance the ability of the clinical laboratory in performing diagnose when culture-independent methods are needed (e.g. in case of bacteremia, septicemia and meningitis), which are rare in the case of campylobacteriosis. The real advantage of implementing WGS in clinical and public heath setting, for what concern *Campylobacter*, is the provision of high-resolution genotyping (including predicting phenotyping) in a faster and, most importantly, automatized way (25). In this contest, species confirmation could function as the first quality assurance step in the analytic pipeline, by using for instance methods especially designed for metagenomics datasets (43) or Average Nucleotide Identity (44).

### Genomic diversity during C. jejuni infection and colonization

The genomic variants that accumulate during human passage should be considered when establishing the level of genetic similarity between epidemiologically linked isolates. The generation of diversity in this population is dependent on the mutation rate and mutational patters of

*C. jejuni*. The fixation of such mutations in the population is again determined by evolutionary processes like genetic drift, bottleneck effect, and selection due to for instance host adaption (45). WGS has been applied to study the genomic diversity and changes accumulating during human infection (46,47) and animal colonization (48,49), of which the latter has been used both as a model organism for human infection [mice; (48,49)] and a reservoir with source attribution [chicken; (49)] in mind. Revez *et al.* (2013) published the first study in which the inoculum isolate (*C. jejuni* NCTC 11168) was subjected to WGS comparison with the excreted isolate-population after human passage (46). The authors demonstrated that the genomic changes occurring during human passage were small and, except for one single nucleotide variant (SNV) in Cj0184c, limited to indels in homopolymeric tracts in contingency loci. These frameshift mutations were generally found in genes regulating surface structures. Bayliss *et al.* (2012) showed that such mutations accumulated rapidly in the population even in the absence of selective pressure, demonstrating mutation rates between ten to hundred times faster than mutation rates in other parts of the genome (45). The same regulatory mechanisms in contingency loci were confirmed in another human-passage study by Thomas *et al.* (2014) (47). Furthermore, the genetic changes observed in *C. jejuni* strains subjected to passage through C57BL/6 IL-10*−/−* mice and IL-10 KO mice were of similar nature: indels in homopolymeric tracts in contingency loci (48,49). However, different loci were affected in mice and chicken relative to humans (48,49), indicating possible differences in adaption mechanisms in animal and human hosts. Thus, care should be taken when extrapolating results from animal colonization experiments to humans. Summarizing, a growing body of evidence points out that phase variation is the major genetic regulatory mechanism during human and animal single passage, enabling *C. jejuni* rapid access to a significant amount of genetic diversity influencing host adaption, virulence and immune evasion by rapid changes in surface structures (45,50).

From an epidemiological standpoint, we propose to exclude such homopolymeric tracts from the WGS comparison in public health operations to minimize the bias introduced in the genetic diversity of the *C. jejuni* strain by the patient itself (20,51–53). In this way, tracing and source attribution of isolates would be independent from genomic changes introduced in the host, and the genomes would more closely resemble that the inoculum isolates.

### Application of WGS of C. jejuni in point-source outbreak investigations

Several studies aim to investigate the applicability of WGS analysis in identifying clonal *C. jejuni* isolates in point-source outbreaks and the retrospective examination of food-, water- and milk-borne outbreaks of campylobacteriosis has gained a lot of research attention. Revez *et al.* (2014) analyzed a resolved milk-borne outbreak of the ST-50 lineage [ST-21 clonal complex (CC)], in which three SNVs and phase variations in 12 loci were identified between the milk- and the patients-isolates by PubMLST-hosted wgMLST-schema featuring 1738 loci (51). The non-related control strains included in the analyses differed by 420-454 alleles from the epidemiologically related strains. In a second study, Revez *et al* (2014) redefined a waterborne outbreak by proving that one of the two investigated cases was a separate, non-related sporadic case; the *C. jejuni* isolated from this patient was deemed too genetically different (69 SNVs) from the assumed source-isolate, while the isolate of the other case differed by only three SNVs compared with the source-isolate (20). In this study, the analysis of homopolymeric tracts was excluded and what was perceived to be homologous recombination was counted as one allele difference (20). Zhang et al. (2015) confirmed the findings of Revez *et al.* (2014a,b) by identifying up to seven allele differences, mainly in homopolymeric runs, between clonal isolates collected from three waterborne outbreaks by the use of an *ad hoc* reference based gene-by-gene approach (Genome profiler; https://sourceforge.net/projects/genomeprofiler/)(54).

Lahti et al. (2016) investigated a foodborne outbreak using WGS in parallel with PFGE and applied a reference-based core genome (cg)-MLST approach using SeqSphere+TM (Ridom© GmbH) to compare the outbreak strains (total of 1271 shared-loci) (55). The authors described a maximum of one allele difference between the clinical isolates. Contrastingly, a *C. jejuni* isolate originating from the same farm as the presumed source (chicken liver paté), but collected 5 months prior to the outbreak, differed by 15 alleles from the patient isolates (55). Furthermore, an Australian study investigated a foodborne outbreak in real-time using WGS and SNVs detection using Snippy v. 3.0 (https://github.com/tseemann/snippy) (56). The study proved the circulation of at least two different STs among patients, forming two distinct clusters. The ST-528 cluster showed SNVs in the range from three to eight, while the ST-535 cluster exhibited 30 SNVs. Moreover, the authors concluded that the source isolate was not “caught” as the *C. jejuni* collected from the suspected source was of a different ST and differed from the case isolates by at least 22,940 SNVs (56). Nevertheless, these results should be interpreted with care as the authors operated with very low coverage (minimum coverage of 10x), they didn’t indicate the overall number of compared nucleotides, and avoid to specify how recombination and homopolymeric tracts were handled (56).

### Application of WGS in surveillance of C. jejuni infections and detection of diffuse outbreaks

The occurrence of diffuse outbreaks, meaning individual cases occurring over a larger area linked to a single vehicle of infection (29), is difficult to detect through routine surveillance, as these cases are masked among the background sporadic cases. However, earlier studies have verified the existence of diffuse outbreaks of campylobacteriosis by demonstrating the presence of temporal, spatial, and genotypic clusters among apparently sporadic cases. Such clusters were two to five times more common than point-source outbreaks in Scotland (29), and accounted for approximately 13% of the reported cases. Similarly, up to 19.6% of the annual campylobacteriosis cases in Norway occurred in spatial-temporal clusters (57). Furthermore, an Alaskan study estimated 32% of the campylobacteriosis cases collected during a decade to be temporal clusters of related PFGE profiles (58). It follows that more emphasis should be put on the detection of such outbreaks, as interventions against the mechanisms creating campylobacteriosis-clusters would be more effective at reducing the burden of illness than actions taken against sporadic transmissions. High-resolution genotyping can improve the detection of clusters of epidemiologically related cases occurring in a less geographically separated group (48) and since WGS offers the highest possible genotyping discrimination, the integration of genome sequencing in surveillance would probably facilitate the implementation of more effective intervention strategies targeted at campylobacteriosis.

Some studies have already integrated genomics in *Campylobacter* surveillance, and all have applied a wgMLST methodology. Cody *et al.* (2013) performed the first real-time genomic epidemiological investigation of *C. jejuni* collected through a surveillance program using a hierarchical wgMLST approach (26,59). In the first all-against-all analysis, BISGdb Genome Comparator, implemented in PubMLST (http://pubmlst.org/), was used to extract the 1026 loci shared among the 379 investigated isolates obtained from the Oxfordshire monitoring program from a total of 1,643 loci listed in the “Gundogdu” schema (http://pubmlst.org/campylobacter), which was established on the re-annotation of *C. jejuni* NCTC 11168 (60). Based on the level of genomic diversity found between isolates collected from a single patient, a cut-off value for investigating the presence of possible clusters was set to 20 allele differences among the 1026 shared-loci (59). Thereafter, an *ad hoc* wgMLST analysis, again using the “Gundogdu schema”, was used to define diversity within the discovered clusters. With this approach, the authors were able to identify temporally associated clusters of strains from cases with apparently no epidemiological links but showing similar genetic diversity as observed between same-patient isolates (3 to 14 loci differences among 1,478 to 1,586 loci shared loci) (59). As observed in point-source outbreaks (20,51,54), the loci that recurrently resulted in differentiation between same-patient and epi-linked isolates were homopolymeric tracts in contingency genes. Similar results were reported by Kovanen *et al.* (2014), who also used a hierarchical wgMLST approach to investigate the genomic relationship of apparently sporadic cases collected during a seasonal peak in Finland (52). First, the authors clustered the isolates based on the 7-loci MLST scheme and performed an *ad hoc* wgMLST analysis using BIGSdb to extract allele information of all 1738 loci for each ST-group. This approach allowed the authors to identify sub-clusters of isolates, which varied only in a few loci, obtained from cases spread over a large geographical area lacking an apparent epidemiological link (52). In their following study, Kovanan *et al.* (2016) attempted to identify the possible source of these predicted diffuse outbreaks, again applying a hierarchical wgMLST approach for each ST group, using the references based gene-by-gene method implemented Genome Profiler (53). Thereafter, the authors manually screened for allele differences, using five SNVs as a cut-off value, while excluding indels in homopolymeric tracts, to define clusters of genetically indistinguishable isolates. The rationale for the low cut-off value was the observed genomic diversity between chicken isolates collected from the same farm during one rearing cycle (61), within point-source outbreaks (20,51,54) and during a human infection (46). When accounting for temporal clustering, the authors were able to link up to 24% of the human cases to specific chicken slaughter batches (53). To identify the possible existence of a diffuse outbreak, Fernandes *et al.* (2015) applied the method introduced by Cody *et al.* (2013) to compared *C. jejuni* isolates collected through a surveillance program with non-related isolates (reference population) of the same ST-21 (62). They showed that 20 of the 23 apparently sporadic cases were indeed part of a cluster, with less than eight (with a mean of four) allele differences across 1577 shared-loci. Simultaneously, the authors demonstrated that the genetic distance between the cluster isolates and the reference population were at least 20 alleles, and proposed faulty pasteurized milk distributed over the investigated catchment area as the source of the clustered cases (62).

The results from these studies support the idea that putative diffuse outbreaks have a clear impact on the epidemiology of campylobacteriosis, and by integrating WGS in the surveillance of *C. jejuni* infections we can potentially distinguish between clustered and sporadic cases. The common belief is that the gene-by-gene method is more suitable for this task than an SNV-calling approach since the gene-by-gene approach is efficient, easy to automate and computationally less intensive.

Although a consensus on the analytic pipeline (experimental and bioinformatical) to use in the gene-by-gene approach and the strain definition cut-off to apply are still under evaluation, it seems clear that the hierarchical wgMLST-pipeline, where a first level genomic clustering based on a core set of loci is performed, followed by an *ad hoc* analysis with an increased number of loci, will be the future reference approach for these types of investigations (59).

## DEFINING THE BASELINE GENOMIC DIVERSITY FOR *C. JEJUNI*

Deciding the genomic diversity present in a population of *C. jejuni* is necessary for deducing whether genomic differences between any two *C. jejuni* isolates are sufficiently small to assume epidemiological relatedness, i.e. the isolates share a common source (3). As seen above, earlier studies have utilized a fairly arbitrary impromptu value for defining epidemiological relationships and genetic associations between strains to detect diffuse outbreaks and trace isolates to possible vehicles and reservoirs. However, the diversity level present within the lineage of which the investigated isolates reside is rarely discussed. Yet, defining such diversity is a necessary prerequisite: if the lineage diversity is generally low, the genetic distinction between two non-linked isolates might fall below the preset cut-off value even when no epidemiological link exists. Evidence proving variation in genetic diversity between *C. jejuni* lineages has recently arisen. For instance, Kovanen *et al.* (2014) found limited genetic diversity within three common STs, namely ST-230, ST-267 and ST-677, while ST-45 were considerably more diverse and separated into three main lineages (52). In a follow-up paper, Llarena *et al.* (2016) investigated the nature of the population structure of ST-45 CC, focusing on identifying possible spatial-temporal evolution signals (21). The authors found that the occurrence and strength of the geographical signal varied between sub-lineages of ST-45 CC strains. Yet, no evidence of a temporal signal was found. In addition, the authors unexpectedly identified certain sub-lineages of ST-45 with extremely similar genomes regardless of time and location of sampling. These successful monomorphic clones were persistently isolated from animal hosts and human patients over a decade from several countries around the globe (21). There is no reason to believe that this clonal nature is limited to ST-45. Indeed, in a recent publication Wu *et al.* (2016) demonstrated the clonal expansion of a highly virulent, ST-8 lineage during the 1970s, which is now causing vast numbers of ovine abortions in the USA (63).

The presence of clonal populations makes genomic distinction between epidemiologically associated and non-related isolates difficult, and seriously hampers the implementation of WGS methodology in surveillance and outbreak investigations for public health purposes. For example, a cluster of ST-45 isolates identified as a diffuse *C. jejuni* outbreak by Kovanen *et al.* (2014) (52) consisted of isolates belonging to a ST-45 population of monomorphic clones (21). Thus, it is uncertain whether these isolates were indeed representatives of a diffuse outbreak, based only on WGS analysis. Clearly, more studies on the genomic diversity and possible occurrence of clonal populations in other *C. jejuni* lineages are needed to resolve this predicament.

## WGS IN *C. JEJUNI* SOURCE ATTRIBUTION AND HOST ADAPTION

Identification of the most frequent transmission routes and foodborne sources of campylobacteriosis is of utmost importance for prioritizing food safety interventions and setting public health goals (64). *C. jejuni* has a complex epidemiology and transmission can occur in numerous ways, including contaminated food, water, and raw milk, and direct animal- and environmental contact. To overcome this problem, methods to quantify the relationship between human patient data and possible infection reservoirs, transmission routes, and risk factors have been developed (65,66). Approaches based on genotypic methods compare the proportion of *C. jejuni* subtypes in different sources and reservoirs with isolates subtypes collected from human patients. These methods rely on different subtypes having variable ability to colonize different hosts or that existing ecological restrictions prevent an equal subtype distribution between reservoirs; often referred to as host adaption (67). Adaptation to a specific host might result in the selection of reservoir-associated traits, such as the presence of specific genes or clusters of genes (68). Therefore, detection of these traits in *C. jejuni* isolates from human patients could theoretically make source attribution simple and straightforward. Although source attribution models using WGS data as input are yet to be applied to campylobacteriosis cases, phylogenic and population structure analyses of *C. jejuni* isolates collected from different studies have been used to trace the source of sporadic cases, as described above (53). On the other hand, major efforts have been devoted to detect and elucidate the mechanisms of host-adaptation in *C. jejuni* with the use of WGS, especially for identifying source-specific traits (67,(69–71). Using 7-loci MLST, Sheppard *et al.* (2011) argued that certain lineages of *C. jejuni* were adapted to wild birds and agricultural animals while other lineages were generalists lacking host adaption (72), and confirmed these findings and attempted to detect the underlying mechanisms of host adaption and generalist lifestyles by WGS and genome-wide association study (GWAS) of *C. jejuni* isolates collected from cattle and chickens (67). The search for host-specific traits within generalist lineages identified a predicted bovine-specific seven-gene region encoding for vitamin B_5_ biosynthesis, and suggested that the overrepresentation of these seven genes in bovines denoted a bacterial adaption to the low vitamin B_5_-containing grass-based bovine-diet. However, the authors pointed out that even though the gene cluster seemed to be necessary in bovines, the fitness costs of such a cluster must have been low as chicken isolates of generalist lineages frequently harboured the same gene region, possibly as a result of a recent host switch from bovines (67). Morley *et al.* (2015) went further in this direction by investigating the molecular mechanisms of host-adaption by analysing one porcine-restricted lineage (ST-403 CC). The authors found three restriction-modification loci unique to this lineage and showed that a number of coding sequences present in other lineages were absent. Moreover, the authors showed that ST-403 CC underwent lineage specific decay and pseudogenization, and they argued that these events were the result of the selective pressure enforced by moving away from avian to porcine host, reflecting the ability of ST-403 CC to differentiate according to niche (71).

The discovery of host-specific loci or gene loss makes these genetic elements possible targets for source tracking. Hence, the genetic make-up of a human isolate could theoretically help determine the source of that isolate. However, we are still far from that possibility, as only a few host-associated loci have been found in *C. jejuni* and the interpretation of their presence is complicated. Additionally, the commonness of generalist-lineages complicates source attribution because of lacking host adaption and signals, probably due to frequent host-jumps (70). Dearlove *et al.* (2015) found that generalists ST-21 CC and ST-45 CC undergo so rapid host jumps that the host signal is totally eroded, and WGS is therefore no more informative than 7-genes MLST in defining the source of infections (70).

## WGS FOR THE DECTION OF *C. JEJUNI* VIRULOME

Differences in strains virulence might explain why certain *C. jejuni* cause more human diseases than others. Consequently, the application of WGS in clinical microbiology might have an advantage in providing a fast and accurate *in silicio* typing (23) of the virulome as defined by several studies. Furthermore, such virulence traits can be used for pinpointing intervention targets for vaccine development. For example, a pan-genome approach led to the discovery of a set of unique loci, or loci with unique alleles, overrepresented in hyper-invasive *C. jejuni* strains which were not related to neutral variation caused by demographic processes (73). Also, bacteremia associated strains showed a linkage between the hyper-invasive phenotype and specific genes within the capsule region (74,75). Moreover, lineages expressing certain lipooligosaccharide structures defined by a specific set of loci are overrepresented in cases of Guillain-Barre syndrome (76,77).

## CONCLUSION

Currently, a few issues are hurdling the full implementation of WGS in real-time surveillance of *C. jejuni* infections: the lack of a common nomenclature and a Tenover-like cut-off value for defining the relationship between *C. jejuni* isolates, and common data analysis tools for enabling comparison of isolates at national and international level. Within these hurdles resides the issue of a limited knowledge on the genetic diversity within different lineages of *C. jejuni,* which impedes the validity of the assumed genetic relationship when WGS is used alone. Additionally, different studies use diverse approaches to compare strains with no clear consensus of which method would be more suitable for meeting the needs and the realities of public health microbiology. Nevertheless, existing evidence supports the benefit of integrating WGS in the surveillance of *C. jejuni* infections and in point-source and diffuse outbreak investigations. Advances in WGS and statistical genetics provide new opportunities to improve our understanding of *C. jejuni* ecology, evolution, and pathogenesis, and have revealed a more complex epidemiological scenario and ecological dynamic than previously believed.

## ACKNOWLEDGEMENT

We thank João André Carriço (University of Lisbon, Portugal) and Alejandra Culebro (University of Helsinki, Finland) for their suggestions for improving the manuscript.

Ann-Katrin Llarena have been supported by INNUENDO project (https://www.innuendoweb.org) co-funded by the European Food Safety Authority (EFSA), grant agreement GP/EFSA/AFSCO/2015/01/CT2 (“New approaches in identifying and characterizing microbial and chemical hazards”).”

## DISCLAIMER

The conclusions, findings, and opinions expressed in this review paper reflect only the view of the authors and not the official position of the European Food Safety Authority (EFSA).

## AUTHORS’ CONTRIBUTIONS

AKL performed the literature search. Both authors contribute in drafting and revising the manuscript. All approved the final draft for publication.

## REFERENCES

1. Köser CU, Ellington MJ, Cartwright EJP, Gillespie SH, Brown NM, Farrington M, et al. Routine Use of Microbial Whole Genome Sequencing in Diagnostic and Public Health Microbiology. PLOS Pathog. 2012 Aug 2;8(8):e1002824.

2. European Food Safety Authority (EFSA). EFSA’s 20th Scientific Colloquium on Whole Genome Sequencing of food-borne pathogens for public health protection. EFSA Support Publ. 2015 Feb 1;12(2).

3. European Centre for Disease Prevention and Control. Expert Opinion on the introduction of next-generation typing methods for food- and waterborne diseases in the EU and EEA. Stockholm: ECDC; 2015.

4. Murray CJL, Vos T, Lozano R, Naghavi M, Flaxman AD, Michaud C, et al. Disability-adjusted life years (DALYs) for 291 diseases and injuries in 21 regions, (1990–2010): a systematic analysis for the Global Burden of Disease Study 2010. The Lancet. 2012 Dec 15;380(9859):2197–223.

5. Humphrey T, O’Brien S, Madsen M. Campylobacters as zoonotic pathogens: A food production perspective. Int J Food Microbiol. 2007;117(3):237–57.

6. Kaakoush NO, Castaño-Rodríguez N, Mitchell HM, Man SM. Global Epidemiology of Campylobacter Infection. Clin Microbiol Rev. 2015 Jul 1;28(3):687–720.

7. Havelaar AH, Pelt W van, Ang CW, Wagenaar JA, Putten JPM van, Gross U, et al. Immunity to Campylobacter: its role in risk assessment and epidemiology. Crit Rev Microbiol. 2009 Feb 1;35(1):1–22.

8. Dingle KE, Colles FM, Wareing DRA, Ure R, Fox AJ, Bolton FE, et al. Multilocus Sequence Typing System for Campylobacter jejuni. J Clin Microbiol. 2001 Jan 1;39(1):14–23.

9. Kärenlampi R, Rautelin H, Schönberg-Norio D, Paulin L, Hänninen M-L. Longitudinal Study of Finnish Campylobacter jejuni and C. coli Isolates from Humans, Using Multilocus Sequence Typing, Including Comparison with Epidemiological Data and Isolates from Poultry and Cattle. Appl Environ Microbiol. 2007 Jan 1;73(1):148–55.

10. Lévesque S, Frost E, Arbeit RD, Michaud S. Multilocus Sequence Typing of Campylobacter jejuni Isolates from Humans, Chickens, Raw Milk, and Environmental Water in Quebec, Canada. J Clin Microbiol. 2008 Oct 1;46(10):3404–11.

11. Sheppard SK, Dallas JF, Strachan NJC, MacRae M, McCarthy ND, Wilson DJ, et al. Campylobacter Genotyping to Determine the Source of Human Infection. Clin Infect Dis. 2009 Apr 15;48(8):1072–8.

12. Strachan NJC, Gormley FJ, Rotariu O, Ogden ID, Miller G, Dunn GM, et al. Attribution of Campylobacter Infections in Northeast Scotland to Specific Sources by Use of Multilocus Sequence Typing. J Infect Dis. 2009 Apr 15;199(8):1205–8.

13. de Haan CPA, Lampén K, Corander J, Hänninen M-L. Multilocus Sequence Types of Environmental Campylobacter jejuni Isolates and their Similarities to those of Human, Poultry and Bovine C. jejuni Isolates. Zoonoses Public Health. 2013 Mar 1;60(2):125–33.

14. de Haan CP, Kivistö RI, Hakkinen M, Corander J, Hänninen M-L. Multilocus sequence types of Finnish bovine Campylobacter jejuni isolates and their attribution to human infections. BMC Microbiol. 2010;10:200.

15. Haan CPA de, Kivistö R, Hakkinen M, Rautelin H, Hänninen ML. Decreasing Trend of Overlapping Multilocus Sequence Types between Human and Chicken Campylobacter jejuni Isolates over a Decade in Finland. Appl Environ Microbiol. 2010 Aug 1;76(15):5228–36.

16. Broman T, Waldenström J, Dahlgren D, Carlsson I, Eliasson I, Olsen B. Diversities and similarities in PFGE profiles of Campylobacter jejuni isolated from migrating birds and humans. J Appl Microbiol. 2004 Apr 1;96(4):834–43.

17. Kärenlampi R, Rautelin H, Hakkinen M, Hänninen M-L. Temporal and Geographical Distribution and Overlap of Penner Heat-Stable Serotypes and Pulsed-Field Gel Electrophoresis Genotypes of Campylobacter jejuni Isolates Collected from Humans and Chickens in Finland during a Seasonal Peak. J Clin Microbiol. 2003 Oct 1;41(10):4870–2.

18. Ragimbeau C, Schneider F, Losch S, Even J, Mossong J. Multilocus Sequence Typing, Pulsed-Field Gel Electrophoresis, and fla Short Variable Region Typing of Clonal Complexes of Campylobacter jejuni Strains of Human, Bovine, and Poultry Origins in Luxembourg. Appl Environ Microbiol. 2008 Dec 15;74(24):7715–22.

19. Magnússon S h., Guðmundsdóttir S, Reynisson E, Rúnarsson Á r., Harðardóttir H, Gunnarson E, et al. Comparison of Campylobacter jejuni isolates from human, food, veterinary and environmental sources in Iceland using PFGE, MLST and fla-SVR sequencing. J Appl Microbiol. 2011 Oct 1;111(4):971–81.

20. Revez M J. Llarena, AK. Schott, T. Kuusi, M. Hakkinen, M. Kivistö, R. Hänninen, ML. Rossi. Genome analysis of Campylobacter jejuni strains isolated from a waterborne outbreak. BMC Genomics. 2014;15(1):1–8.

21. Llarena A-K, Zhang M J. Halkilahti, J. Hänninen, ML. Rossi, Vehkala M, Välimäki N, Hakkinen M, Hänninen M-L, et al. Monomorphic genotypes within a generalist lineage of Campylobacter jejuni show signs of global dispersion. Microb Genomics. 2016 Sep 14;Ahead of Print.

22. Biggs PJ, Fearnhead P, Hotter G, Mohan V, Collins-Emerson J, Kwan E, et al. Whole-Genome Comparison of Two Campylobacter jejuni Isolates of the Same Sequence Type Reveals Multiple Loci of Different Ancestral Lineage. PLOS ONE. 2011 Nov 11;6(11):e27121.

23. Carrillo CD, Kruczkiewicz P, Mutschall S, Tudor A, Clark C, Taboada EN. A framework for assessing the concordance of molecular typing methods and the true strain phylogeny of Campylobacter jejuni and C. coli using draft genome sequence data. Front Cell Infect Microbiol. 2012;2:57.

24. Köser CU, Holden MTG, Ellington MJ, Cartwright EJP, Brown NM, Ogilvy-Stuart AL, et al. Rapid Whole-Genome Sequencing for Investigation of a Neonatal MRSA Outbreak. N Engl J Med. 2012 Jun 14;366(24):2267–75.

25. Didelot X, Bowden R, Wilson DJ, Peto TEA, Crook DW. Transforming clinical microbiology with bacterial genome sequencing. Nat Rev Genet. 2012 Sep;13(9):601–12.

26. Jolley KA, Maiden MC. BIGSdb: Scalable analysis of bacterial genome variation at the population level. BMC Bioinformatics. 2010;11:595.

27. Sheppard SK, Jolley KA, Maiden MCJ. A Gene-By-Gene Approach to Bacterial Population Genomics: Whole Genome MLST of Campylobacter. Genes. 2012 Apr 12;3(2):261–77.

28. Centers for Disease Control and Prevention (CDC) Office of Workforce and Career Devlopment. Epidemic disease occurrence. In: CDC, editor. Principles of Epidemiology in Public Health Practice - An Introduction to Applied Epidemiology and Biostatistics [Internet]. 3rd ed. Atlanta, GA, USA: CDC; 2012. p. 1–72. Available from: http://www.cdc.gov/ophss/csels/dsepd/ss1978/lesson1/section11.html

29. Strachan NJC, Forbes KJ. Extensive Spatial and Temporal Clustering of Campylobacter Infections Evident in High-resolution Genotypes. In: Sheppard SK, Meric G, editors. Campylobacter ecology and evolution. 1st ed. Norfolk, UK: Caister Academic Press; 2014. p. 253–63.

30. European Commission. Commission implementing decision of 8 August 2012 amending Decision 2002/253/EC laying down case definitions for reporting communicable diseases to the Community network under Decision No 2119/98/EC of the European Parliament and of the Council (notified under document C(2012 5538). Off J Eur Union. 2012;(262).

31. Hänninen M-L, Perko-Mäkelä P, Rautelin H, Duim B, Wagenaar JA. Genomic Relatedness within Five Common Finnish Campylobacter jejuni Pulsed-Field Gel Electrophoresis Genotypes Studied by Amplified Fragment Length Polymorphism Analysis, Ribotyping, and Serotyping. Appl Environ Microbiol. 2001 Apr 1;67(4):1581–6.

32. Tenover FC, Arbeit RD, Goering RV, Mickelsen PA, Murray BE, Persing DH, et al. Interpreting chromosomal DNA restriction patterns produced by pulsed-field gel electrophoresis: criteria for bacterial strain typing. J Clin Microbiol. 1995 Sep;33(9):2233–9.

33. Taboada EN, MacKinnon JM, Luebbert CC, Gannon VP, Nash JH, Rahn K. Comparative genomic assessment of Multi-Locus Sequence Typing: rapid accumulation of genomic heterogeneity among clonal isolates of Campylobacter jejuni. BMC Evol Biol. 2008;8:229.

34. Skirrow MB. Campylobacter enteritis: a “new” disease. Br Med J. 1977 Jul 2;2(6078):9–11.

35. Butzler JP, De Boeck M, Goossens H. New selective medium for isolation of Campylobacter jejuni from faecal specimens. Lancet Lond Engl. 1983 Apr 9;1(8328):818.

36. Totten PA, Patton CM, Tenover FC, Barrett TJ, Stamm WE, Steigerwalt AG, et al. Prevalence and characterization of hippurate-negative Campylobacter jejuni in King County, Washington. J Clin Microbiol. 1987 Sep;25(9):1747–52.

37. Butzler J-P. Campylobacter, from obscurity to celebrity. Clin Microbiol Infect. 2004 Oct;10(10):868–76.

38. Denis M, Soumet C, Rivoal K, Ermel G, Blivet D, Salvat G, et al. Development of a m-PCR assay for simultaneous identification of Campylobacter jejuni and C. coli. Lett Appl Microbiol. 1999 Dec 1;29(6):406–10.

39. Kärenlampi RI, Tolvanen TP, Hänninen M-L. Phylogenetic Analysis and PCR-Restriction Fragment Length Polymorphism Identification of Campylobacter Species Based on Partial groEL Gene Sequences. J Clin Microbiol. 2004 Dec;42(12):5731–8.

40. Korczak BM, Stieber R, Emler S, Burnens AP, Frey J, Kuhnert P. Genetic relatedness within the genus Campylobacter inferred from rpoB sequences. Int J Syst Evol Microbiol. 2006 May;56(Pt 5):937–45.

41. Murray PR. Matrix-assisted laser desorption ionization time-of-flight mass spectrometry: usefulness for taxonomy and epidemiology. Clin Microbiol Infect. 2010 Nov;16(11):1626–30.

42. Bessède E, Solecki O, Sifré E, Labadi L, Mégraud F. Identification of Campylobacter species and related organisms by matrix assisted laser desorption ionization–time of flight (MALDITOF) mass spectrometry. Clin Microbiol Infect. 2011 Nov;17(11):1735–9.

43. Sankar A, Malone B, Bayliss SC, Pascoe B, Méric G, Hitchings MD, et al. Bayesian identification of bacterial strains from sequencing data. Microb Genomics [Internet]. 2016 [cited 2016 Sep 26];2(8). Available from: http://mgen.microbiologyresearch.org/content/journal/mgen/10.1099/mgen.0.000075

44. Konstantinidis KT, Tiedje JM. Genomic insights that advance the species definition for prokaryotes. Proc Natl Acad Sci U S A. 2005 Feb 15;102(7):2567–72.

45. Bayliss CD, Bidmos FA, Anjum A, Manchev VT, Richards RL, Grossier J-P, et al. Phase variable genes of Campylobacter jejuni exhibit high mutation rates and specific mutational patterns but mutability is not the major determinant of population structure during host colonization. Nucleic Acids Res. 2012 Jul 1;40(13):5876–89.

46. Revez M-L J. Schott, T. Llarena, AK. Rossi, M. Hänninen. Genetic heterogeneity of Campylobacter jejuni NCTC 11168 upon human infection. Infect Genet Evol. 2013;16:305–9.

47. Thomas DK, Lone AG, Selinger LB, Taboada EN, Uwiera RRE, Abbott DW, et al. Comparative Variation within the Genome of Campylobacter jejuni NCTC 11168 in Human and Murine Hosts. PLOS ONE. 2014 Feb 7;9(2):e88229.

48. Jerome JP, Bell JA, Plovanich-Jones AE, Barrick JE, Brown CT, Mansfield LS. Standing Genetic Variation in Contingency Loci Drives the Rapid Adaptation of Campylobacter jejuni to a Novel Host. PLOS ONE. 2011 Jan 24;6(1):e16399.

49. Kim J-S, Artymovich KA, Hall DF, Smith EJ, Fulton R, Bell J, et al. Passage of Campylobacter jejuni through the chicken reservoir or mice promotes phase variation in contingency genes Cj0045 and Cj0170 that strongly associates with colonization and disease in a mouse model. Microbiology. 2012;158(5):1304–16.

50. Parkhill J, Wren BW, Mungall K, Ketley JM, Churcher C, Basham D, et al. The genome sequence of the food-borne pathogen Campylobacter jejuni reveals hypervariable sequences. Nature. 2000 Feb 10;403(6770):665–8.

51. Revez M-L J. Zhang, J. Schott, T. Kivistö, R. Rossi, M. Hänninen. Genomic variation between Campylobacter jejuni isolates associated with milk-borne-disease outbreaks. J Clin Microbiol. 2014;52(8):2782–6.

52. Kovanen M-L SM. Kivistö, RI. Rossi, M. Schott, T. Kärkkäinen, UM. Tuuminen, T. Uksila, J. Rautelin, H. Hänninen. Multilocus sequence typing (MLST) and whole-genome MLST of campylobacter jejuni isolates from human infections in three districts during a seasonal peak in Finland. J Clin Microbiol. 2014;52(12):4147–54.

53. Kovanen M-L S. Kivistö, R. Llarena, AK. Zhang, J. Kärkkäinen, UM. Tuuminen, T. Uksila, J. Hakkinen, M. Rossi, M. Hänninen. Tracing isolates from domestic human Campylobacter jejuni infections to chicken slaughter batches and swimming water using whole-genome multilocus sequence typing. Int J Food Microbiol. 2016;226:53–60.

54. Zhang M J. Halkilahti, J. Hänninen, ML. Rossi. Refinement of whole-genome multilocus sequence typing analysis by addressing gene paralogy. J Clin Microbiol. 2015;53(5):1765–7.

55. Lahti E, Löfdahl M, Ågren J, Hansson I, Olsson Engvall E. Confirmation of a Campylobacteriosis Outbreak Associated with Chicken Liver Pâté Using PFGE and WGS. Zoonoses Public Health. 2016 Jun 1;n/a-n/a.

56. Moffatt CRM, Greig A, Valcanis M, Gao W, Seemann T, Howden BP, et al. A large outbreak of Campylobacter jejuni infection in a university college caused by chicken liver pâté, Australia, 2013. Epidemiol Amp Infect. 2016 Jun;1–8.

57. Jonsson ME, Heier BT, Norström M, Hofshagen M. Analysis of simultaneous space-time clusters of Campylobacter spp. in humans and in broiler flocks using a multiple dataset approach. Int J Health Geogr. 2010;9:48.

58. Castrodale LJ, Provo GM, Xavier CM, McLAUGHLIN JB. Calling all Campy – how routine investigation and molecular characterization impacts the understanding of campylobacteriosis epidemiology – Alaska, United States, (2004–2013). Epidemiol Amp Infect. 2016 Jan;144(2):265–7.

59. Cody AJ, McCarthy ND, Rensburg MJ van, Isinkaye T, Bentley SD, Parkhill J, et al. Real-Time Genomic Epidemiological Evaluation of Human Campylobacter Isolates by Use of Whole-Genome Multilocus Sequence Typing. J Clin Microbiol. 2013 Aug 1;51(8):2526–34.

60. Gundogdu O, Bentley SD, Holden MT, Parkhill J, Dorrell N, Wren BW. Re-annotation and reanalysis of the Campylobacter jejuni NCTC11168 genome sequence. BMC Genomics. 2007;8:162.

61. Llarena A-K, Huneau A, Hakkinen M, Hänninen M-L. Predominant Campylobacter jejuni Sequence Types Persist in Finnish Chicken Production. PLOS ONE. 2015 Feb 20;10(2):e0116585.

62. Fernandes AM, Balasegaram S, Willis C, Wimalarathna HML, Maiden MC, McCarthy ND. Partial Failure of Milk Pasteurization as a Risk for the Transmission of Campylobacter From Cattle to Humans. Clin Infect Dis Off Publ Infect Dis Soc Am. 2015 Sep 15;61(6):903–9.

63. Wu Z, Periaswamy B, Sahin O, Yaeger M, Plummer P, Zhai W, et al. Point mutations in the major outer membrane protein drive hypervirulence of a rapidly expanding clone of Campylobacter jejuni. Proc Natl Acad Sci. 2016 Sep 6;201605869.

64. Pires SM, Evers EG, van Pelt W, Ayers T, Scallan E, Angulo FJ, et al. Attributing the human disease burden of foodborne infections to specific sources. Foodborne Pathog Dis. 2009 May;6(4):417–24.

65. Hald T, Vose D, Wegener HC, Koupeev T. A Bayesian approach to quantify the contribution of animal-food sources to human salmonellosis. Risk Anal Off Publ Soc Risk Anal. 2004 Feb;24(1):255–69.

66. Mullner P, Spencer SEF, Wilson DJ, Jones G, Noble AD, Midwinter AC, et al. Assigning the source of human campylobacteriosis in New Zealand: a comparative genetic and epidemiological approach. Infect Genet Evol J Mol Epidemiol Evol Genet Infect Dis. 2009 Dec;9(6):1311–9.

67. Sheppard SK, Didelot X, Meric G, Torralbo A, Jolley KA, Kelly DJ, et al. Genome-wide association study identifies vitamin B5 biosynthesis as a host specificity factor in Campylobacter. Proc Natl Acad Sci. 2013 Jul 16;110(29):11923–7.

68. Champion OL, Gaunt MW, Gundogdu O, Elmi A, Witney AA, Hinds J, et al. Comparative phylogenomics of the food-borne pathogen Campylobacter jejuni reveals genetic markers predictive of infection source. Proc Natl Acad Sci U S A. 2005 Nov 1;102(44):16043–8.

69. Baily JL, Méric G, Bayliss S, Foster G, Moss SE, Watson E, et al. Evidence of land-sea transfer of the zoonotic pathogen Campylobacter to a wildlife marine sentinel species. Mol Ecol. 2015 Jan 1;24(1):208–21.

70. Dearlove BL, Cody AJ, Pascoe B, Méric G, Wilson DJ, Sheppard SK. Rapid host switching in generalist Campylobacter strains erodes the signal for tracing human infections. ISME J. 2016 Mar;10(3):721–9.

71. Morley L, McNally A, Paszkiewicz K, Corander J, Méric G, Sheppard SK, et al. Gene Loss and Lineage-Specific Restriction-Modification Systems Associated with Niche Differentiation in the Campylobacter jejuni Sequence Type 403 Clonal Complex. Appl Environ Microbiol. 2015 Jun;81(11):3641–7.

72. Sheppard SK, Colles FM, McCARTHY ND, Strachan NJC, Ogden ID, Forbes KJ, et al. Niche segregation and genetic structure of Campylobacter jejuni populations from wild and agricultural host species. Mol Ecol. 2011 Aug 1;20(16):3484–90.

73. Baig A, McNally A, Dunn S, Paszkiewicz KH, Corander J, Manning G. Genetic import and phenotype specific alleles associated with hyper-invasion in Campylobacter jejuni. BMC Genomics. 2015;16:852.

74. Kivistö M-L RI. Kovanen, S. Skarp-De Haan, A. Schott, T. Rahkio, M. Rossi, M. Hänninen. Evolutionand comparative genomics of Campylobacter jejuni ST-677 clonal complex. Genome Biol Evol. 2014;6(9):2424–38.

75. Skarp CPA, Akinrinade O, Nilsson AJE, Ellström P, Myllykangas S, Rautelin H. Comparative genomics and genome biology of invasive Campylobacter jejuni. Sci Rep. 2015 Nov 25;5:17300.

76. Revez J, Hänninen M-L. Lipooligosaccharide locus classes are associated with certain Campylobacter jejuni multilocus sequence types. Eur J Clin Microbiol Infect Dis. 2012 Feb 2;31(9):2203–9.

77. Taboada EN, van Belkum A, Yuki N, Acedillo RR, Godschalk PC, Koga M, et al. Comparative genomic analysis of Campylobacter jejuni associated with Guillain-Barré and Miller Fisher syndromes: neuropathogenic and enteritis-associated isolates can share high levels of genomic similarity. BMC Genomics. 2007;8:359.

